# Personalizing computational models to construct medical digital twins

**DOI:** 10.1101/2024.05.31.596692

**Authors:** Adam C. Knapp, Daniel A. Cruz, Borna Mehrad, Reinhard C. Laubenbacher

**Author notes:** A.C.K and D.A.C. contributed equally to this work. B.M. and R.C.L. conceived of the project design and guided the research. All authors contributed to the conceptualization of the project. A.C.K. and D.A.C. developed, validated, and applied the mathematical algorithms; algorithms were implemented by A.C.K. All authors contributed to the writing of the manuscript. The authors declare no competing interest.

## Abstract

Digital twin technology, pioneered for engineering applications, is being adapted to biomedicine and healthcare; however, several problems need to be solved in the process. One major problem is that of dynamically calibrating a computational model to an individual patient, using data collected from that patient over time. This kind of calibration is crucial for improving model-based forecasts and realizing personalized medicine. The underlying computational model often focuses on a particular part of human biology, combines different modeling paradigms at different scales, and is both stochastic and spatially heterogeneous. A commonly used modeling framework is that of an agent-based model, a computational model for simulating autonomous agents such as cells, which captures how system-level properties are affected by local interactions. There are no standard personalization methods that can be readily applied to such models. The key challenge for any such algorithm is to bridge the gap between the clinically measurable quantities (the macrostate) and the fine-grained data at different physiological scales which are required to run the model (the microstate). In this paper we develop an algorithm which applies a classic data assimilation technique, the ensemble Kalman filter, at the macrostate level. We then link the Kalman update at the macrostate level to an update at the microstate level that produces microstates which are not only compatible with desired macrostates but also highly likely with respect to model dynamics.

**Significance Statement:** To realize the promise of personalized medicine, we need to be able to integrate different types of data collected from a given patient into a computational framework that enables decision making about optimal interventions to help this patient to either maintain or regain health. Digital twins represent such a framework, once the technology is sufficiently developed. A fundamental problem that currently does not have a widely applicable solution is how to calibrate a generic computational model of human biology to a given patient at a given time. This paper presents a solution to this problem for the agent-based model framework, commonly used to capture stochastic and spatially heterogeneous biological processes, such as tumor growth or immune system dynamics.

---

**A** medical digital twin (MDT) is a virtual replica of some aspect of a patient’s biology relevant to their health; it consists of a computational model that is calibrated to the patient and is dynamically updated from data repeatedly collected from the patient. The MDT can be used to forecast a patient’s health trajectory or simulate the effect of different interventions. MDTs promise to be a key technology for the implementation of personalized medicine. While there are some examples that deliver on this promise, such as the artificial pancreas (1), several foundational problems need to be solved first (2, 3) before this technology can be used widely.

In this paper, we address the core mathematical problem of calibrating a generic computational model to a particular patient, forecasting disease progression, and the effect of therapeutic interventions. We study data assimilation (DA), the integration of observational data into a computational model, for the purpose of personalizing the model. Specifically, we consider how to find likely states and parameterizations of the model so that we can improve the prediction of a patient’s health outcomes. Finding the precise state and parameters of a patient, especially given small amounts of available data, is a challenging task: models may have structurally or practically unidentifiable parameters in this regime (4), and many model states and parameters may reproduce the observed data. To tackle this challenge, we employ parameter and state distributions that capture these uncertainties.

One of the features that sets applications of DT technology to human health apart from other areas, such as numerical weather prediction or most industrial applications, is that there are different model types for different aspects of human biology. In particular, many biological features are characterized by spatial heterogeneity, such as lung biology, or by their inherently stochastic nature, such as the immune system. This makes it often infeasible to use the framework of ordinary differential equations, for which many mathematical tools are available. An increasingly important modeling framework in this context are agent-based models (ABMs), a mechanistic modeling framework which is common in cellular biology (5–8) and biomedicine, with applications ranging from respiratory diseases to immune-mediated diseases and cancer (9–15), and is supported by a growing list of software platforms (16–19). For instance, elements of the immune response to pathogens can be effectively captured with this framework by taking advantage of spatial heterogeneity to properly capture the effect of small numbers of cells in certain regions, such as resident macrophages in lung alveoli.

Detailed mechanistic models are capable of reproducing a variety of patient outcomes (see (1, 20, 21)) and are motivated by the need to optimize mechanism-based interventions. A drawback is that, in addition to clinically measurable quantities, they often include practically unobservable quantities. These mostly relate to the basic feature of ABMs that the individual agents (e.g., cells) have internal states, spatial positions, and interact with each other and the spatial environment, generating what we refer to as “microstates” which contain the fine-grained information required to compute the model. In contrast, measurements taken from a patient that can be used to personalize a generic mechanistic model to the patient, typically capture “macrostates” – aggregate measures such as blood cytokine levels or lung immune cell counts. The interplay between micro- and macrostates lies at the heart of the DA challenges for ABMs, and is the topic of the current manuscript.

Extensive work has been done on applying DA techniques to such ABMs (22–24), especially in the context of transportation systems (25–35) and social/epidemiological spread (36–42); however, little work has been done related to biomedical applications. Moreover, the existing research generally falls into two categories: (i) DA done using full microstates of ABMs and (ii) DA done on surrogate models built from an ABM. For the former, considering the full microstate of an ABM will typically include granular details which are not medically relevant, will likely rely on data that may not be available, and may take on a form which is incompatible with standard techniques. The high-dimensionality of such a system requires a very large ensemble to give adequate estimates of many state variables, resulting in high computational costs inappropriate to the better-than-real-time requirements of an MDT. For the latter, MDTs will require the inclusion of mechanistic controls in the underlying model to allow physicians to evaluate the effects of therapies. Designing surrogate models that include such mechanistic controls is a challenging task in its own right which has not been resolved generally (43, 44).

The mathematical problem we face is the following: We are given an ABM, together with repeated partial measurements of macrostates. With each repeated measurement, we need to recalibrate the ABM and simulate it to make a prediction about its likely trajectory. For this purpose, we will use an ensemble Kalman filter method (45) on macrostates, used extensively in other contexts. These macrostate modifications need to be translated to equivalent microstate modifications, in order to generate new ABM predictions. Aside from an application of Kalman filter methods to ABMs in a biomedical context, the main technical contribution of this paper is a microstate synthesis algorithm that generates the “most likely” microstates associated to a given macrostate.

We apply the general algorithm that we developed to two example ABMs. The first is a spatially well-mixed model of predator-prey dynamics (46) for which the microstate synthesis problem is relatively simple. For this model we get good performance of the algorithm overall. However, we find that there are certain cases that the present algorithm cannot handle. These occur for certain absorbing states of the model that cannot be distinguished by the observables at hand. The second is a model of viral dynamics (47) which exhibits a number of more complex features in the microstate, including a high degree of spatial heterogeneity. To deal with this increase in complexity, we apply a microstate synthesis algorithm which employs quantization and error diffusion. For this model, our analysis uncovered the existence of several model phenotypes and we explored the performance of the algorithm at the point where these phenotypes diverge.

## Background

The process of integrating patient data into a computational model is an example of *data assimilation* (DA); the study of how to optimally combine observed data and theory, in the form of a mathematical or computational model, into a predictive forecast. In order to provide a concrete example of the type of eventual applications we envision for our algorithm, we describe a scenario below in an Intensive Care Unit setting where a doctor might benefit from a decision support tool. Such a tool would provide forecasts of the patient’s health trajectory, based on mechanistic simulation, and possible interventions available to the doctor.

### Remark 1

*We present the case of a patient cared for by one of the authors (BM) as a prototypical scenario to which simulation-based decision support algorithms could be applied: The patient was a 41 year old man with class I obesity (body mass index 34) and no other medical history who presented to the Emergency Department in November 2021 after a 9 day history of fever, cough, myalgia, and progressive shortness of breath. On presentation, he was febrile, had normal blood pressure, and had an oxygen saturation on room air was 85%. His chest x-ray showed bilateral airspace disease, and he tested positive for COVID-19. He required endotracheal intubation and mechanical ventilation for acute respiratory distress syndrome, and was treated with remdisivir and dexamethasone. Over the next 12 days, he developed worsening ARDS, septic shock requiring escalating doses of vasopressors, and acute kidney injury. He died on the 13th hospital day of refractory shock*.

In the context of an MDT, one of our principal goals is to determine what kinds of interventions can be applied to a patient like the one in Remark 1 in order to improve outcomes. This requires that the model is sufficiently well calibrated, or *personalized*, to the patient to give accurate predictions of trajectories and the effects of interventions. However, realistic mechanistic models typically have high dimensional and complex states, far exceeding the amount of data that can be expected from clinically feasible patient measurements. After model construction, the next task in building an MDT is to develop techniques to solve the ill-posed inverse problem of estimating the patient’s current state from the available data.

Typical methods for solving ill-posed inverse problems include forms of regularized maximum likelihood optimization, such as Tikhonov regularization, or are based on Bayesian statistical methods. It is not clear how regularization alone could provide a quantification of uncertainty into its forecasts, a key component of MDTs (2). For this reason, we have opted to use a Bayesian filtering technique to predict patient outcomes, inspired by Kalman filter (KF) methods used in numerical weather forecasting (48–54).

Bayesian frameworks have several advantages, including a natural means of using prior knowledge of reasonable initial conditions and parameter values from a reference population. The use of an informative prior is an important technique to deal with the large number of parameters in complex models and paucity of data that can be collected clinically, especially early in treatment. Bayesian techniques also provide a convenient context with which to integrate uncertainty into forecasts. In particular, KF methods have several moving parts which are described statistically: 1) a predictive model, 2) a true/target state, and 3) observations/measurements. These methods have two steps:

1. Predict: Given an initial distribution for states at time *t*_*n*_, the model is used to produce predictive state distributions for times *T > t*_*n*_.
2. Update: When a measurement is taken at time *t*_*n*+1_ *> t*_*n*_, the predictive distribution at time *t*_*n*+1_ is updated based on the measurement and predictive distribution at *t*_*n*+1_.

The updated distribution at time *t*_*n*+1_ is then fed back into the predictive step as the initial distribution of states at time *t*_*n*+1_ and the process begins again. An initial state distribution at *t* = 0 is required to start the process.

The Kalman filter (55–57) is the closed form solution to the Bayesian filtering equations in the case where the dynamic and measurement models are linear stochastic and all distributions are (multivariate) Gaussian. We discuss the filter in more detail in the Supplementary Information; however, we note that there have been numerous extensions of the KF including the extended Kalman filter (EKF) (58, 59), ensemble Kalman filter (EnKF) (45, 60, 61) and unscented Kalman filter (UnScKF) (62), each of which contains tunable hyper-parameters related to measurement and process uncertainty. In this work, we employ the EnKF because it tends to be most suitable for nonlinear stochastic models and is often at the core of several numerical weather prediction models (49, 50, 52–54, 61). The linear stochastic dynamic model of the KF is replaced by a general computational model of dynamics in the EnKF, and the predictive distributions 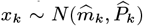 are computed by selecting an *ensemble* of samples from *N*(*m*_*k−*1_, *P*_*k−*1_), running the computational model on each in order to obtain an ensemble of samples for the predictive distribution.

An appropriate measure is necessary to evaluate KF performance. Average or relative distances between ensemble mean predictions and the corresponding true trajectory do not account for predictive uncertainty. We chose a concept from information theory called *surprisal* (63) which takes these uncertainties into account by measuring the amount of information gained from revealing the true trajectory given a predictive distribution; see the Supplementary Information for definitions. Informally, surprisal indicates how surprised we should be to learn the true trajectory; thus, high probability events will have low surprisal and vice versa. Finally, the KF includes hyper-parameters related to measurement uncertainty and process uncertainty which need to be adapted to the disease model at hand.

## Results

### The Kalman Filter on ABMs

In order to make use of KF methods in agent-based (or other) models with complex state spaces, we will need to take care to define several of the mathematical objects considered in the sketch above. First, it is useful to consider which state components to include, not limiting ourselves to quantities for whom the model includes dynamics, but also including quantities that are often thought of as model parameters, as long as they vary between individuals and have a meaningful impact on model dynamics. As discussed below, is is often useful to expand the model to include random walk dynamics on these quantities in order to improve KF performance. Further, the complex state spaces of agent-based models lead to several complications, which lead us to a distinction between fully-described “microstates” and more summarized “macrostates” upon which the KF acts. See Figure 1(a)

**Fig. 1.**
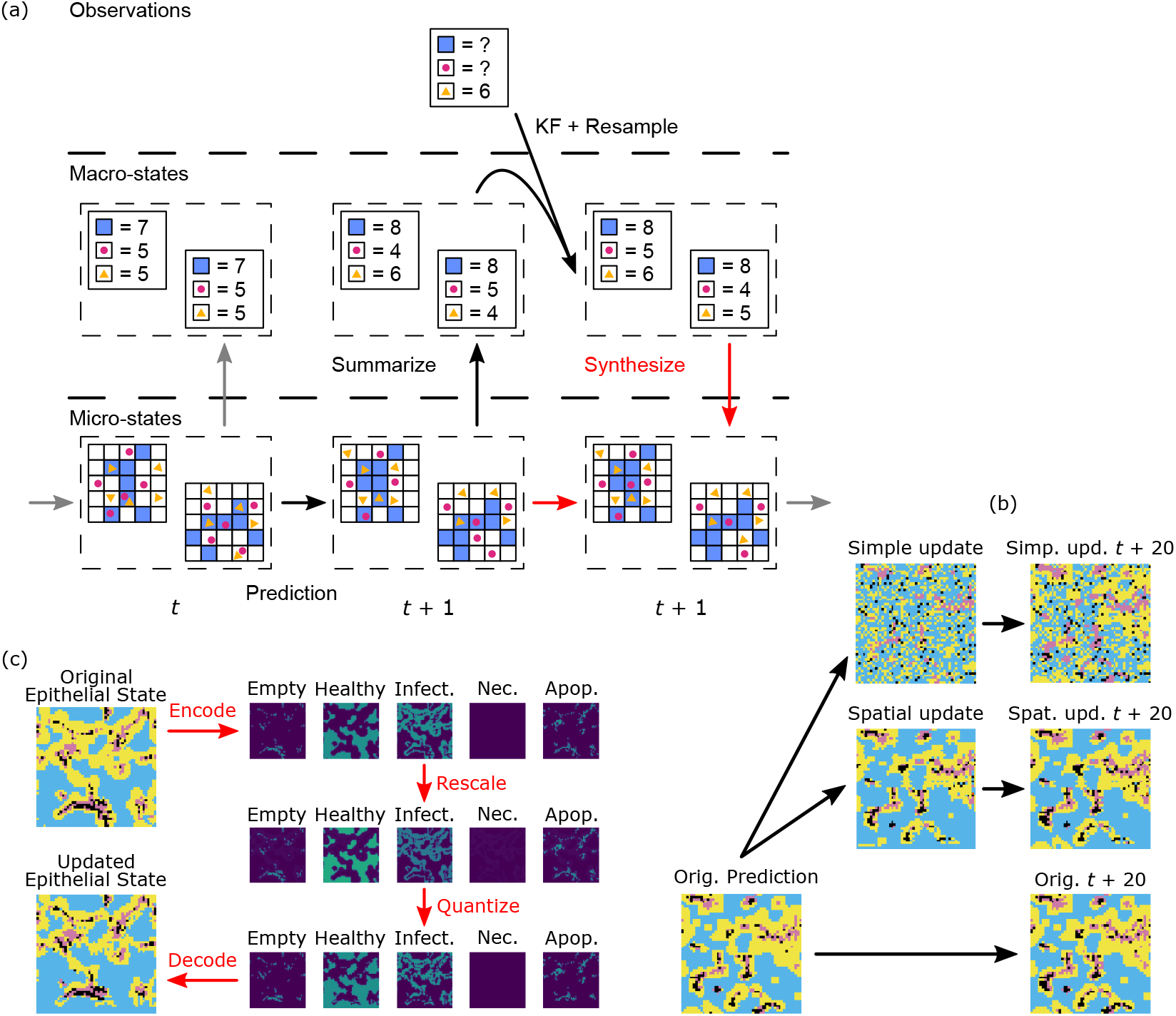
(a) Overview of the ABM Kalman Filter (ABMKF) algorithm’s update of microstates based on updates to the macrostates. Prediction: An ensemble of microstates at time *t* is advanced to *t* + 1 using the mechanistic model. Summarize: The updated microstates are then summarized into macrostates. Resulting macrostates are used as the ensemble in the Ensemble Kalman filter (EnKF). Update: The EnKF resamples the updated ensemble of macrostates. Synthesize (in red): The previous microstates are used as seeds for the synthesis of new macrostate-compatible microstates. (b) Starting epithelial microstate (time *t*), Second column: microstate updated by the two synthesis algorithms, Third column: microstates 20 time steps in the future. (c) Categorical epithelial states are encoded into five component scalar states as indicator variables. This “one-hot encoding” (64) is rescaled to increase the healthy and decrease the infected component sums. The scaled field is quantized back to a one-hot encoding using the quantization/error-diffusion algorithm. Finally, the quantized one-hot encoding is decoded as a categorical state.

While we will not attempt to provide a formal definition of an ABM, we note that many computational models that go by this name have some commonalities. (7, 8, 65–67) Such a model contains one or more types of agent. For example, in the Wolf-Sheep-Grass (WSG) model considered later in the Results, wolves and sheep are mobile agents and each grass patch is an immobile agent. An ABM of the immune system might contain several agent types, one for each type of immune cell considered as well as epithelial and endothelial cells, such as in the viral dynamics model discussed in later in the Results. Such an ABM will typically support multiple instances of each agent type, each of which has its own state described by the same collection of data and updated by the same computational model. Often, these agents may enter or leave the simulation through a birth/death process.

These features lead to problems with applying KF methods directly to such a model:

1. The space of states for the ABM may be broken into components which vary in dimension. The birth/death process for agents changes the total amount of data required to completely describe the model state, leading to dimensional changes when the number of agents change. This means that an ensemble of model instances may have members whose states do not live in the same dimension.
2. There is often no natural matching of agents between ensemble members. This means that, even in the absence of a birth/death process, there is no *a priori* way of identifying the state spaces of model instances.
3. An agent’s state may not be a vector space. For example, an ABM may drive agent behavior through a Boolean network or a categorical state, leading to a (partially) discrete state space. Since the KF naturally produces continuous vector valued states, models would not be able to use these directly and require the development of discretization techniques.
4. Many ABMs include spatial components leading to excessively high dimensionality: a molecular density field might correspond to a variable taking values over an *N × M* lattice. That is, it corresponds to *N × M* dimensions in the global state space. Given *K* such molecules, we have at least *KMN* dimensions in the global state space and the covariance matrices used in the KF would then have up to 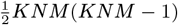 entries to estimate. For example, in the viral dynamics model considered below we have *K* = 12, *M* = *N* = 51 so that we are discussing the estimation of over 487 million values in the covariance matrix. Given that the likely observables of such fields are merely the *K* spatial averages, this is a lot of computational cost to bear.

A possible solution to both problems is suggested by considering analogous problems in statistical mechanics and kinetics. In statistical mechanics, one makes a distinction between microstates and macrostates. A *microstate* is a complete description of a system’s state. For a gas, this might be a list of the position and momentum of all its constituent particles. In this setting, a *macrostate* is a description of the gas as a whole such as pressure, volume, number of molecules, and temperature.

We intend to perform data assimilation using KF methods on macrostates, as implied in (40) and fully proposed in (41). However, there is an immediate problem: our models require the full microstate to run, and while the KF gives us a method of updating the distribution of model macrostates after an observation, it does not immediately lead to methods whereby we can update the corresponding distribution of microstates.

In our context, we propose some possible components to define the macrostate of the model so that it is useful for KF methods:

- For a collection of *N* agents, a quantity that is invariant under agent permutation. e.g., in a gas, the sum of the momenta. Averages and (co)variances of agent properties.
- The number of agents of a given type, especially when that number is relatively large.
- Average or total values of spatially distributed quantities. Small neighborhood statistics.
- Realistically measurable quantities. For MDTs, this may include measurement of various substances measured in blood but exclude measurements with detailed spatial information about the distribution of these substances in an organ.

These quantities have some desirable qualities such as being relatively stable over short times, frequently more compatible with realistic measurements, and being approximately continuous.

While the summarization map from microstates to macrostates is a straightforward computation, the map is many-to-one. This means that, given a macrostate, there are many possible microstates representing it and the *microstate synthesis* step in Figure 1(a) requires some choices. In the statistical mechanics analogy, we know that for a given macrostate (*P, T, n, V*) there are many possible ways of arranging the *n* particles in 3-space. In statistical mechanics, a fundamental assumption is that each of these configurations is equally likely. This assumption may or may not be true for any given agent-type in an ABM and we will see examples of both below. When the equiprobable assumption is true, there is a clear algorithm to synthesize microstates: generate a random microstate compatible with the macrostate through some uniform procedure. This could be generalized to the case where there is an explicit expression of microstate distributions for each macrostate, but it is very common that such an expression will be unavailable. We describe one technique for this case of microstate synthesis in the Results section below.

The purpose of the MDT will determine a minimum of which components must included in the macrostate. In principle, all likely clinical measurements which could be derived from the model and its microstates could be included in the macrostate. However, these come at a cost, as the number of samples required to accurately estimate the means and covariances in the KF depends on the overall dimension of the macrostate space. We select the macrovariables based on a combination of measures reported in the source articles as well as expert input as to what could be clinically measurable. An application of this method should consider trade-offs between the information gained by including a macrovariable and the additional computational cost. Beyond the inclusion or omission of a macrovariable, we must also consider variable transformations. See the Supplementary Information for more details.

### Microstate Synthesis for Biological ABMs

Extending KF based methods to ABMs requires a microstate synthesis algorithm. We propose a general principle for microstate synthesis algorithms for EnKFs and specific implementations for the models we present as use cases. Constructing such an algorithm has been straightforward in prior work (40, 41) because of the presence of discrete model compartments which house agents which correspond to a comparatively small difference between micro and macrostates; these compartments allow for the macrostate to be composed of only agent counts per compartment. However, the presence of such compartments is not guaranteed in general, especially in ABMs which feature spatial attributes.

The first step for our algorithm is initialization of the ensemble. These initial microstates will depend on parameters sampled from a Gaussian prior distribution on macrostates. For the examples of interest here, it is comparatively straightforward to create the details of the corresponding initial microstates as they will be in a known configuration. e.g., in the patient described in the introduction, an estimate of their state at the onset of infection. This initialization gives us an initial micro- and macrostate ensemble.

The prediction/correction, which we call the ABMKF, algorithm is then:

1. Advance the microstate ensemble using the model, producing samples from a predictive microstate distribution.
2. Summarize predictive microstate trajectories to predictive macrostate trajectories.
3. Using a macrostate-derived measurement, perform ensemble KF methods to the predictive macrostate trajectories to obtain a Gaussian posterior distribution for macrostates.
4. Sample a new ensemble of macrostates from the posterior distribution obtained from the EnKF.
5. For each member of the new macrostate ensemble synthesize a compatible microstate.

A diagram of this procedure is shown in Figure 1(a). Step 5 requires additional explanation. How do we generate these microstates? Both intuition and our physics analogy suggest that whatever microstate *μ* we choose to represent the macrostate *M* should be one which is realistic in the sense that it is highly likely for the model to be in that state. That is, we would like to choose *μ* based on the probability distribution *p*(*μ* | *M*). For simple models, it may be possible to describe all or parts of *p*(*μ* | *M*) explicitly, e.g., agent positions and velocities in the Wolf-Sheep-Grass model. In that case, we can generate *μ* by sampling from said distribution. However, as the complexity of the model increases, our ability to describe *p*(*μ* | *M*) explicitly is reduced and we must develop other methods.

The methods for microstate synthesis that we present here are based on the following principles and assumptions:

- Continuity: If *M*_0_ and *M*_1_ are nearby macrostates, then the distributions *p*(*μ* | *M*_0_) and *p*(*μ* | *M*_1_) are similar. That is, if *μ*_0_ is a microstate for *M*_0_, then there is a microstate *μ*_1_ for *M*_1_ with a similar likelihood and which is close to *μ*_0_. See Figure 1(b).
- Local similarity: In spatial models, agents in similar local neighborhoods should have similar trajectories.

The continuity assumption suggests the following algorithm: Given a collection 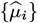 of predictive microstates and 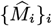 the corresponding collection of macrostates and *{M*_*i*_*}* a sample from the KF posterior, pick a pairing 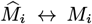 which minimizes pairwise change. (We use the Gail-Shapely algorithm.) The continuity assumption then tells us that there should be microstates *{μ*_*i*_*}* for the macrostates *{M*_*i*_*}* that are pairwise close to the 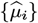. Thus, the problem becomes: how do we synthesize a microstate *μ*_*i*_ for *M*_*i*_, given a similar microstate 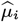 which has a similar macrostate 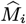.

By passing to the macrostate, we resolve the first and second problems associated to applying KF methods to ABMs. Recall the third of our problems: that an agent’s state may not be a vector space. We propose a generalizable solution for agents taking a categorical state and implement it, and a corresponding microstate synthesis algorithm which respects non-uniform spatial distributions, for epithelial cells in the viral dynamics model. This approach can be applied to models in which agents positioned on a grid can take one of several categorical states.

For the fourth problem, the dimensionality of spatial components, we note that naïvely viewing a density field as a vector exploits none of the spatial relations of the model such as similarity between local neighborhoods or the possible spatial symmetries of the model. Both the WSG and viral dynamics models are invariant under affine transformations of the plane. For this reason, we enhance our quantization-error diffusion algorithm to also impute molecular field levels based on local data. For details, see the section on Quantization and Error Diffusion in the Supplementary Information.

### Spatially well-mixed Predator-Prey Dynamics

Our first example is a relatively simple ABM of predator-prey dynamics, the Wolf-Sheep predation model (46, 68, 69), a member of the Netlogo (70) model library which we have re-implemented in python (71) under the name of the Wolf-Sheep-Grass (WSG) model. In this model, wolves, sheep, and grass exist in a two-dimensional space. Grass occupies fixed patches in this space while wolves and sheep move though space in a random walk. Each grass pixel can be either on or off, indicating the presence of grass, and wolves and sheep keep an internal energy counter. Wolves, resp. sheep, eat sheep, resp. grass, at their location, each gaining energy in the process, and grass regrows over time. Wolves and sheep both reproduce randomly, with some probability, splitting their energy between the parent and child. See Table 1 for a summary of the variables present in the model.

**Table 1.**
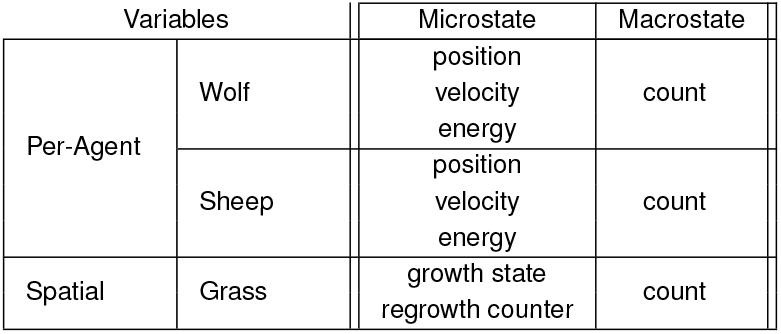
Micro- and macro-state variables for the WSG model. Each wolf and sheep agent has a position, velocity and energy while grass agents are described by their growth state (green/brown) and regrowth timer. Macrostate summaries of these agents are given by the count of wolf and sheep agents and the count of grass agents in the alive state.

As seen in Table 1, we choose to describe a macrostate of this model in terms of the number of wolves, sheep, and proportion of grass coverage. As both wolves and sheep move in a random walk, their observed spatial distributions are approximately uniform, as is the distribution of grass. This suggests that, once a macrostate has been specified, it is straightforward to construct a likely spatial distribution of wolves, sheep and grass by uniform random sampling.

As described above, KF methods require some degree of tuning in order to achieve good performance. With the components of the macrostate chosen, the next type of tuning to consider consists of variable transformations. In the WSG model, it is relatively common for wolves to go extinct and so the support for the wolf distribution is often concentrated around zero. For this reason, we make a variable transformation on the number of wolves, *W* ^*′*^ = log *W*. Note that this itself adds a certain degree of complication, since the number of wolves can actually be zero. We resolve this by using a slight variant of the log transform, instead working with *W* ^*′*^ = log(*ϵ* + *W*) and the inverse transform *W* = max(0.0, exp(*W* ^*′*^) − *ϵ*) where *ϵ* = 0.001.

As the KF is a minimum least squares estimator, the KF tends to learn the largest magnitude quantities very strongly while treating other interesting, but lower magnitude, parameters more like noise. This means that variable transformations, such as scaling, may also be required to achieve good performance of the KF. Notably, the number of sheep and grass are typically one and two orders of magnitude more than the number (or log number) of wolves, respectively, while the parameters tend to be around the same order of magnitude as the log-wolves. For this reason, we scale the number of sheep by 0.1 and the grass count by 0.01, respectively, to keep these variables in similar ranges.

Measurement uncertainty was tuned by simulation with the following procedure. For each choice of measurement and measurement uncertainty, we generate a population of virtual “true” trajectories from the model. That is, we generate a sample of random parameterizations of the WSG model and run each through 1000 time steps to form a population of true trajectories. We then run the KF methods described above, with the chosen measurement taken at regular intervals, and evaluate the data-assimilation process using surprisal. With 1000 samples each, we evaluated the three basic measurements (i.e. wolves, sheep, or grass) and four levels of measurement uncertainty (i.e. 10.0, 1.0, 0.1, and 0.01). The results are shown in Figure 2a. Note that the lowest level tested generally, but not always, performed best.

**Fig. 2.**
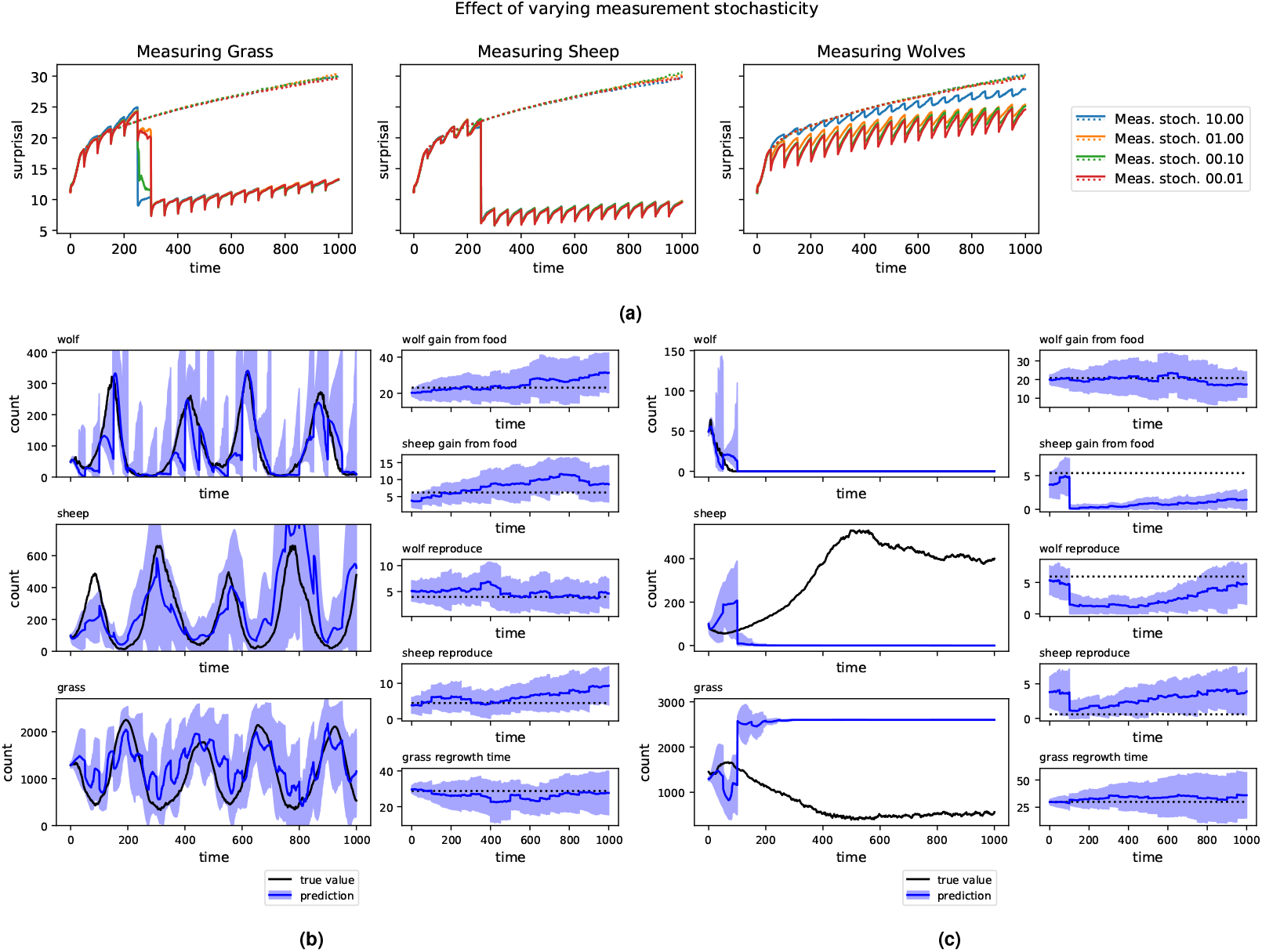
A: Median surprisal over 1000 runs as a time series for the original ensemble and after a series of KF update steps. Measurements taken on wolves, sheep, and grass, respectively. Downward spikes in surprisal correspond to KF updates. *Q* = 0.01. Spatial dims=255 *×* 255. B: A typical run of the ABMKF algorithm where measurements are taken on the count of wolves. In black, the true trajectory. In blue, the distribution of trajectories learned, with uncertainty cone. The corrections to the wolf count are close to the true value with a corresponding narrowing of the uncertainty bands after measurement. The amount of narrowing and degree to which the mean moves to the observation depend on the measurement uncertainty. The KF similarly corrects the sheep and grass counts reasonably well, despite these counts not being directly measured. As such, the uncertainty bands around these predictions do not change as much. C: A very high surprisal trajectory. In black, the true trajectory. In blue, the distribution of trajectories learned, with uncertainty cone. In the true trajectory, wolves have gone extinct and are the sole measurement being taken. In this system, there are two possible equilibria consistent with wolf-extinction: one where the sheep stabilize around a high level, and another where the sheep are also extinct. In this case, the KF stabilized around the extinct-sheep fixed point, but the system is near the high-sheep attractor. This incorrect choice cannot be corrected by further wolf measurements.

Process stochasticity was also tuned by simulation in a similar manner. Note that here, we only add process stochasticity to the parameters, as the computational model includes inherent stochasticity for the state variables. Also, the added stochasticity is only added to the model instances used in the KF; the parameters of the true trajectory are kept constant. See Figure S1.

In Figures 2b and 2c, we show examples of trajectories learned. In Figure 2b, measurements are taken on the number of sheep and we see a relatively good example of the KF correcting errors in prediction. While the KF incorrectly predicts that wolves will go extinct, leading to upward wandering predictions on the number of sheep, measurements on the number of sheep keep the overall prediction from completely diverging from the true trajectory. However, Figure 2c shows a failure mode for KF methods. In this example, the KF is attempting to learn from periodic measurements of wolves, but in fact wolves have gone extinct. While the KF successfully learns that the number of wolves is zero, it strongly misidentifies the state of sheep and grass. That is, the wolf-free model has two attractor states: one which is a random walk around a large number of sheep and small quantity of grass and another in which sheep are extinct and grass is at a steady state. In Figure 2c, the random chance of sampling yields the incorrect option and, due to the unimodal nature of Gaussian distributions, gets stuck there. This example serves to demonstrate the importance of measurement type as either additional sheep or grass measurements could distinguish between these attractors.

### Spatially Heterogeneous Viral Dynamics

The second ABM which we will examine is a model of human anti-viral immune responses(47) whose basic structure is relevant to the ICU patient described earlier. This model is significantly more complex than the WSG model, both in the number of agents and parameters as well as in the behavior of the agents. In this model, spatial distributions of agents are distinctly non-uniform and play a significant role in the model’s evolution.

Briefly, the model is spatial, on a two-dimensional lattice of size 51 *×* 51. It contains several immune cells as mobile agents: dendritic cells, pulmonary neutrophils, macrophages, and natural killer cells. Fixed at each lattice point, the model includes agents modeling the epithelium and endothelium as well as the quantity of extracellular virus and various molecular species. The human portion of the model has approximately 60 parameters and 42 microstate variables which partially correspond to 20 macrostate variables; see Table S1. A sensitivity analysis selects 19 of the 60 parameters; see the Supplementary Information for more detail.

The dimension of a microstate for this model is sizable from the perspective of data assimilation, with 20 variables at each of the 51 *×* 51 lattice sites. Each of the mobile agents contributes at least a location (2D) and movement direction (2D) to the microstate as well as other fields depending on the type of agent. In our Python implementation of this model, on a typical 64-bit computer, the microstate takes around 0.95 megabytes of memory, depending somewhat on the number of agents. This puts us in the regime where it is possible to hold many instances of the model in memory simultaneously, but where inference from low-dimensional data is challenging.

Among the possible macrostate components, we consider several which could potentially be measurable using non-destructive procedures such as a bronchoalveolar lavage (BAL) or through blood work. In particular, we select 22 quantities to describe the macrostate of the model: total quantities of all modeled molecules, amounts of extracellular and intracellular virus, counts of lung epithelial and endothelial cells in various states, counts of the various immune cell agents, and a count of the number of apoptotic epithelial cells which have been killed by the virus. For this introductory study, we measure these values directly from the simulation.

In the WSG model, the distributions of wolves, sheep, and grass were close to uniform and we were able to reconstruct compatible microstates by simple random sampling. However, in the viral dynamics model, spatial distributions of cytokines and cells are highly non-uniform and are characterized by “hot-spots” where the infection is present. This complicates the synthesis of compatible microstates. We developed and tested two microstate synthesis algorithms. The first is a simple update technique similar to that which was used for the WSG model. The second is a more complex, spatially aware technique that makes modifications that try to maximize the similarity of local neighborhoods to previously seen local neighborhoods.

For some model quantities, there are simple methods to update microstates. For example, in the case of a molecular field, such as `IL1`, an update to the `total IL1` macrostate can be propagated to the `IL1` microstate by appropriate scaling. This method preserves spatial correlations.

However, this method does not work for all model quantities. For example, consider the update of the counts of epithelial cells. In this model, epithelial cells are located at patches and every patch in the simulation can take several categorical states in which the patch is empty or contains a healthy, an infected, or dead epithelial cell. Dead epithelial cells are distinguished by process: apoptosis or necrosis.

The macrostate update may change the total number of cells in each category but the total number in all categories remains constant and equal to the number of patches. In the simpler of the microstate synthesis algorithms, we change the microstate to the desired macrostate by altering the states of randomly selected cells. This achieves a certain type of minimality to the microstate update: it minimizes the total number of changed epithelial states.

Since this random alteration is not chosen with regard to the geometry of the infection, we have no expectation that this manner of update will preserve the hot spots. In fact, we typically see that this form of update will tend to create new hot-spots as newly placed infected epithelial cells will often be outside of existing infected regions. See Figure 1(b). To remedy this problem, we introduced an update technique which better preserves the existing geometry of a microstate. Consider the case of the categorical state of epithelial cells in the viral dynamics model. The existing technique of scaling, which we performed for molecular concentrations, cannot be done directly on a categorical state. Instead we developed a 3-step process: 1) encode each cell’s categorical state as a vector, through what is known as one-hot encoding in machine learning (64) or as a dummy-variable in statistics, 2) scale the resulting vector field to be compatible with the desired macrostate, and 3) quantize the vector field back to a categorical state while preserving the global counts of the various categories. In more detail,

1. *One-hot encoding of the previous state*. Each patch’s state takes values in one out of five categories: empty, or containing a healthy, infected, necrotic, or apoptosed epithelial cell. The one-hot encoding is then a mapping of these categorical states to the standard basis vectors of R^5^: Healthy → *e*_1_, Infected → *e*_2_, Necrosed → *e*_3_, Apoptosed → *e*_4_, Empty → *e*_5_. Note that the counts of each type are now the components of the vector obtained by summing the one-hot encoded vectors. See Figure 1(c).
2. *Rescaling*. Given the initial epithelial-state count macrostate (*H, I, D, A, E*) and updated epithelial-state count macrostate (*H*^*′*^, *I*^*′*^, *D*^*′*^, *A*^*′*^, *E*^*′*^), we rescale the one-hot vectors by component-wise multiplication by (*H*^*′*^*/H, I*^*′*^*/I, D*^*′*^*/D, A*^*′*^*/A, E*^*′*^*/E*). These new vectors are no longer one-hot encoded, but do have the property that the sum of sites will have the updated macrostate, (*H*^*′*^, *I*^*′*^, *D*^*′*^, *A*^*′*^, *E*^*′*^). See Figure 1(c)
3. *Quantization and error diffusion*. The goal of this step is to return the epithelial vectors to a one-hot encoding while preserving the updated macrostate. Our procedure for this involves two steps: First, based on the state vector 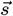 of each epithelial cell and possibly its surroundings, we must choose an appropriate *quantization* to one of the standard basis vectors *e*_*i*_. These algorithms preserve various qualitative and quantitative properties and have an almost hundred-year-long history in image processing (72, 73). Instead of optimizing quantization for visual perception, we will base the quantization on the minimum of a multi-component “loss function” which balances various targets:

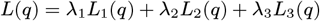

where *L*_*i*_(*q*) are target functions related to quantization error, neighborhood structure, and neighborhood likelihood, respectively, and *λ*_*i*_ are scalar weights. See the section on Quantization and Error Diffusion in the Supplementary Information for an explicit definition of each target function. This quantization introduces an error term, 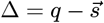, to the macrostate. This error is then uniformly *diffused* to any surrounding unquantized epithelial cells (see Figure S2). By distributing the error to surrounding epithelial cells, we preserve the overall macrostate.

We repeat this 3-step process for the endothelial cells as well, which have three categorical states: Normal, Activated, and Dead.

As stated above, there are multiple options for how to appropriately choose a quantization and multiple options for the pattern of error diffusion, each of which can be arbitrarily paired in the overall quantization/error-diffusion algorithm. We consider here an alternative to choosing the closest quantization, which incorporates data from the local neighborhood. We seek a quantization that seeks to minimize a weighted combination of three things:

1. The quantization error, represented by 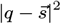. This influences the quantization to be close to 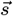.
2. The dissimilarity of the quantization to neighbors, represented by the negative logarithm of *c · q*, where *c* is a regularized count of the number of each type of cell in a 3 *×* 3 neighborhood. This influences the quantization to be similar to neighboring values, reducing the number of isolated types.
3. The negative log probability of the neighborhood based on previous runs. This measure, which is based on both local cells states and local cytokine levels, influences the algorithm to pick quantizations that have been previously observed.

See the Supplementary Information for further details.

Initial tuning of the KF for the viral dynamics model revealed interesting effects. Notably, there is a spike in uncertainty from around *t* = 500 to *t* = 750 when tuning the measurement uncertainty and a similar but longer lasting spike when tuning the process stochasticity. In each case, the spike in uncertainty is resolved after further time and measurements. See Figure 3a for measurement stochasticity and Figure S3 for process stochasticity.

**Fig. 3.**
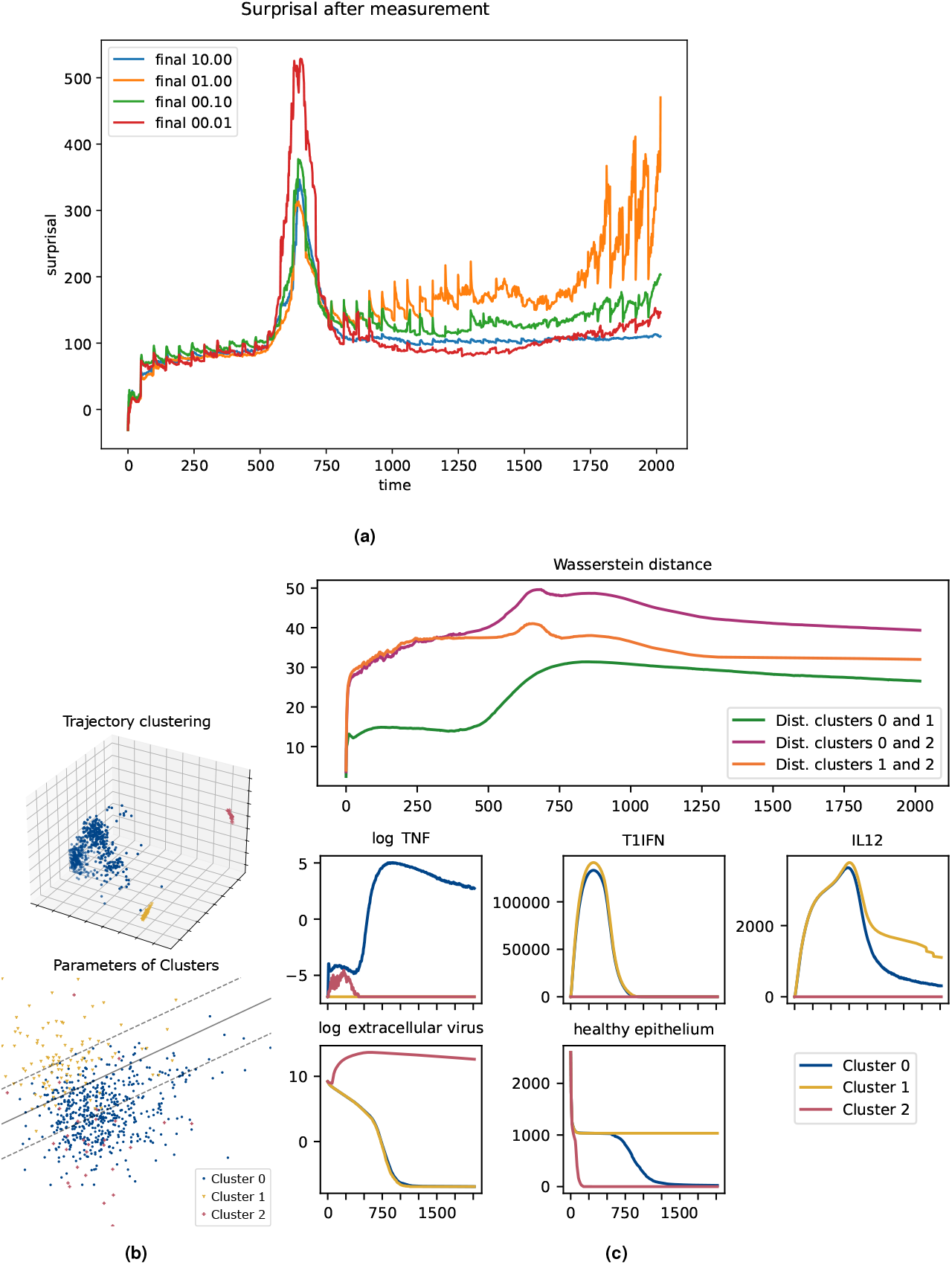
(a) Tuning the measurement uncertainty for the viral dynamics model. In the spike from *t* = 500 to *t* = 750, the curve with the lowest measurement uncertainty has the highest surprisal due to overfitting by the ABMKF during a period of divergence in trajectories. (b) Dimensional reduction of trajectories using PCA shows clusters that suggest phenotypes. Left: trajectories, suggesting three clusters, corresponding to possible phenotypes. Right: parameters, labeled by cluster, showing approximate linear boundary between clusters 0 and 1. Separating lines computed using `scikit-learn`’s `LinearSVC`. (c) Top plot: Time series of Wasserstein distances between clusters. At each time-point the Gaussian approximation of each cluster is calculated and the pair-wise Wasserstein distances are calculated between the resulting three Gaussians. Cluster 2 distinguishes itself rapidly from the others, while clusters 0 and 1 have an abrupt separation between *t* = 500 and *t* = 750. Lower rows shows a representative selection of macrostate components. TNF (shown in log levels), P/DAMPS, IL1, IL6, and IL10 are present at low levels in both clusters 0 and 1 until the clusters diverge around *t* = 500. Similarly, healthy epithelial counts also distinguish these clusters at around this time. Clusters 0 and 1 have similar trajectories in T1IFN (shown) and IFNg for the entire course of the infection, while these clusters have divergent IL12 (shown) and IL8 levels in the later stage of the infection. Cluster 2 is distinguished by extremely high levels of virus (shown in log levels), and rapid epithelial cell death.

To explore what led to this phenomenon, we collated the virtual patient trajectories generated while running the tuning process. This resulted in 695 simulation runs consisting of 41 variables sampled at 2017 time points. These were flattened into 695 samples in a 2017 *×* 41 dimensional space and principal component analysis (PCA) was used to dimensionally reduce the vectors into 3 dimensions. See the left subfigure in Figure 3b. This reduction showed that the trajectories fall into at least 3 meaningfully distinct clusters, suggesting a possible decomposition of the trajectories into phenotypes. To evaluate the possibility that these conjectured phenotypes are driven by model parameterization, we performed PCA on our 695 samples to reduce the 19-dimensional model parameterizations to 3 dimensions. The first dimension of this PCA was found to be an uninformative artifact of rounding certain parameters to integer values. The second and third dimension are used to plot the left subfigure in Figure 3b. This 2-dimensional plot shows a rough, but distinct, linear separation pattern between clusters 0 and 1 which corresponds to a separating hyperplane in the full 19-dimensional parameter space. This confirms that these clusters are in part driven by model parameterizations. Note that cluster 2, having only 28 representatives, does not have a clear pattern with respect to this hyperplane.

In Figure 3c, we see some justification for the idea that these clusters drive the spike in surprisal and correspond to a divergence between phenotypes at that time. Consider the top subfigure in Figure 3c in which we see the time-wise change between each cluster in terms of Wasserstein distance (74), a well-established measure of the difference between probability distributions. Cluster 2 separates quickly from the others, which can be seen from the rapid growth and sustained height of the orange and purple curves. The distance between clusters 0 and 1, however, remains comparatively low until around *t* = 500, transitioning to a sustained higher level by *t* = 750. This change should be interpreted as a divergence in the phenotypes in that region. In effect, we see three phases for the surprisal: First, a lower surprisal phase where the phenotypes are not very distinct. Second, a higher surprisal phase, where the phenotypes diverge and the KF is uncertain which corresponds to the virtual patient. Finally, a third phase which returns to lower surprisal levels as the KF has resolved the virtual patient’s phenotype. This interpretation is backed up by examining the mean cluster trajectories, as seen in the lower plots of Figure 3c.

This clustering phenomenon, corresponding to the divergence of phenotypes, brings up an interesting point. As noted in (75), the KF is an optimal minimum mean square error estimator (MMSEE), regardless of the underlying distribution type. In that article, the main thrust is that the optimality of the KF does not depend on the Gaussian assumption and that the KF is then an optimal choice in more situations than commonly believed. However, the diverging phenotypes seen above are exactly the sort of phenomenon in which an MMSEE is not a reasonable estimate. This type of estimator is most appropriate for unimodal data; when there are multiple clusters, an MMSEE will often predict something in the low-likelihood region in-between the clusters.

## Discussion

In this work, we introduced a microstate synthesis algorithm that allows us to apply KF methods to agent-based models. As a first application, we applied this algorithm to a predatorprey model. Given that the model was spatially well-mixed, we chose a simple approach for microstate synthesis in which agent generation and removal actions were carried out using uniform random distributions. With this approach, our algorithm performed well, with notable exceptions occurring when virtual patients and/or ensemble members ran into boundary states where one or more agent types went extinct. We then applied our data assimilation algorithm to a viral dynamics model. Because the spatial heterogeneity of this model strongly influences system dynamics, we opted for a more spatially aware approach to microstate synthesis. This spatially aware microstate synthesis algorithm, which carries out the continuous Kalman update in discrete settings using a quantization algorithm, performed well overall. At the same time, the discovery of a spike in surprisal revealed how the KF performs when the model contains several phenotypes which diverge mid-simulation.

An important direction for our future work is to study the general applicability of this personalization method and required modifications or extensions beyond the two use cases we presented here. One major challenge is that there is great diversity in the structure, features, and implementations of agent-based models, and an absence of standards for their specification and implementation. Another challenge involves modeling a measurement procedure itself, such as the location of blood draw and analytic device used (76–78). As the two use cases show, even for relatively simple models, application and analysis of our algorithms is complex. In part, we will address this issue by focusing on models implemented in one of the standard modeling platforms, including CompuCell3D (16), TissueForge (79), PhysiCell (80), and others. This introduces a certain standardization of model structure and opens the possibility of standardizing at least part of our algorithm.

In other future work, we will introduce a technique to deal with the kind of no-exit sets that were observed in the WSG model. This extension, which we have given the preliminary name of “Stratified Kalman Filter”, partitions the state space into strata based on the no-exit sets, keeping track of the likelihood that the model is in each stratum while performing KF techniques within each stratum. We will also work on improving microstate synthesis using modern AI techniques such as generative adversarial networks. Additionally, we will consider techniques to better deal with phenotype-based non-linearities in systems such as seen in the viral dynamics model. As a diagnostic, the detection of phenotypes provides valuable qualitative information for a physician. However, as qualitative measures, phenotypes have different properties from either the continuous-numerical macrostates or from microstates; these differences require further investigation and modeling. We expect that, when further expanding the applications of the KF to stochastic *multiscale* models, such as a model (9, 81) our laboratory developed to study the immune response to respiratory fungal infections all of the above issues will persist. Such models will require further new techniques which can bridge the various scales.

As we have discussed above, the data assimilation tools developed in this paper could serve as one component of a digital twin. Such a twin can serve as a virtual experimental setup where a physician can test possible medical interventions against the predicted effects on the personalized digital twin. This human-in-the-loop setup can be expanded to consider actions other than strictly therapeutic treatments, including the question of which measurements to take. That is, the model could provide guidance as to which measurements will be most informative at any given time, allowing the digital twin to advise physicians how proposed measurements will affect predictions and either order or skip certain diagnostics. Tools have been developed in control theory (82, 83) to answer exactly these measurement questions, but mostly targeted to the case where there is no human in the loop and the decision of what to measure is answered once and for all. In future work, we will explore the human-in-the-loop aspect of measurement and the question of how best to offer guidance to the user on all of the available actions.

## Data and Code Availability

Implementation details, code, and data are freely available on GitHub under the Apache 2.0 license at: https://github.com/LaboratoryForSystemsMedicine/enkf-on-abms

## ACKNOWLEDGMENTS

A.C.K. was supported by NSF Grant 2325776. B.M. was supported by awards NIH 1 R01 HL169974-01, NIH 1 R011AI135128-01, NIH 1 R01 HL169974-01. R.C.L. acknowledges funding from the following awards: U.S. Army ACC-APG-RTP W911NF, NIH 1 R01 HL169974-01, U.S. DoD DARPA HR00112220038, NIH 1 R011AI135128-01, NIH 1 R01 HL169974-01.

