## Supplementary Material for "Personalizing computational models to construct medical digital twins"

#### This PDF file includes:

Supporting text
Figs. S1 to S6
Tables S1 to S5
Legends for Movies S1 to S2
SI References

#### Other supporting materials for this manuscript include the following:

Movies S1 to S2

### Supporting Information Text

**The Kalman Filter.** In a Bayesian filter, we hope to find the likely distribution of the system state $x_k$ at time $t = k$, conditional on observations $y_1, \ldots, y_k$ at the current and previous time points. The Kalman filter (KF) is the closed form solution to the Bayesian filtering equations in the case where the dynamic and measurement models are linear stochastic and all distributions are multivariate Gaussian. These assumptions reduce the update algorithm to linear algebra. That is:

$$x_k = A_{k-1}x_{k-1} + q_{k-1} \qquad y_k = H_k x_k + r_k$$

where $A_{k-1}$ is the dynamic model, $q_{k-1} \sim N(0, Q_{k-1})$ is the process noise, $H_k$ is the measurement model, and $r_k \sim N(0, R_k)$ is the measurement noise. Under these assumptions, the prediction step becomes:

$$\widehat{m}_k = A_{k-1}m_{k-1} \qquad \widehat{P}_k = A_{k-1}P_{k-1}A_{k-1}^T + Q_{k-1}$$

where $m_k, P_k$ are the parameters of the posterior state distribution $x_k \sim N(m_k, P_k)$ conditional on the observation $y_k$ and $\widehat{m}_k, \widehat{P}_k$ are the parameters of the predicted state distribution $\widehat{x}_k \sim N(\widehat{m}_k, \widehat{P}_k)$. The update step becomes:

$$v_k = y_k - H_k\widehat{m}_k \qquad S_k = H_k\widehat{P}_kH_k^T + R_k \qquad K_k = \widehat{P}_kH_k^TS_k^{-1}$$
$$m_k = \widehat{m}_k + K_kv_k \qquad P_k = \widehat{P}_k - K_kS_kK_k^T$$

where $v_k$ is the prediction error and $K_k$ is called the Kalman gain matrix. The KF is further the unique *optimal linear* minimum mean squared error estimator (1) for $x_k$, and the Gaussian-distribution assumptions are not required (2) for optimality.

In the Ensemble Kalman Filter (EnKF) (3, 4), we instead approximate the machinery above by sampling. We start with an initial ensemble $\widehat{x}_0^{(i)}$ with mean $\mu_0$ and covariance $\Sigma_0$. These ensemble members can then be advanced to any desired $k$ using simulation to ensembles $\widehat{x}_k^{(i)}$ at which the corresponding statistics $\widehat{\mu}_k$, $\widehat{\Sigma}_k$ can be computed. Then, when a measurement $y_k$ is taken at time $k$, the Kalman update can be performed exactly as before, as the sensitivity and Kalman-gain matrices are computed using only the prediction $\widehat{\mu}_k$, the measurement $y_k$, $\widehat{\Sigma}_k$, the measurement matrix $H_k$, and the measurement uncertainty matrix $R_k$.

**Surprisal.** To evaluate the prediction performance, we employ the Shannon information or surprisal (ref). Given a probability density function $p$, the surprisal of the event $X = x$ is $-\log(p(X = x))$. Thus, the surprisal of the event $X = x$ under the Gaussian distribution $N(\mu, P)$ is

$$\frac{1}{2}(x-\mu)^T\Sigma^{-1}(x-\mu) + \frac{1}{2}\log\det\Sigma + \frac{D}{2}\log 2\pi$$

where $D$ is the dimension of the (state) space associated with $N(\mu, \Sigma)$. Surprisal is closely related to the Mahalanobis distance for normal distributions, which is defined by the quadratic form part of the surprisal: $(\gamma(t) - \widehat{\mu}(t))^T\widehat{\Sigma}^{-1}(t)(\gamma(t) - \widehat{\mu}(t))$.

From the previous section, recall that KF methods produce Gaussian-valued predictive trajectories or equivalently a map $\widehat{\gamma} : [t_n, t_{n+1}] \to \mathbb{R}^D \times \mathbb{R}^{D\times D}$ defined by $\widehat{\gamma}(t) \to (\widehat{\mu}(t), \widehat{\Sigma}(t))$. The average surprisal of the real trajectory under the predictive distribution is then:

$$\frac{1}{2(t_{n+1} - t_n)}\int_{t_n}^{t_{n+1}} \left[(\gamma(t) - \widehat{\mu}(t))^T\widehat{\Sigma}^{-1}(t)(\gamma(t) - \widehat{\mu}(t)) + \log\det\widehat{\Sigma}(t) + D\log 2\pi\right] dt$$

**Macrostate Variable Transformations.** Many variables and parameters of interest in a biological model are naturally non-negative and KF methods are based on Gaussian distributions. When the distribution of a macrovariable tends to stay away from zero, we may work with approximately Gaussian distributions such as $|X|$ or $\max(0, X)$ where $X$ is Gaussian distributed. However, when a macrovariable tends to spend a lot of time at or near zero, it may be worthwhile to transform the variable. e.g., via a log-transform. Further, as with many computational algorithms based on best $L^2$-fits, the KF tends to perform better when the various macrovariables are reweighted to be on similar scales.

In the wolf-sheep-grass model, we used the Kalman filter on macrostate variables which were

- transformed by $w \mapsto \log(10^{-3} + w)$ for the number of wolves,
- divided by 10 for the number or sheep, and
- divided by 100 for the count of grass.

All other state variables (parameters) were unchanged.

In the An-Cockrell model, all macrostate variables were transformed by the function $\log(10^{-3} + x)$.

**Implementation Details for Quantization and Error Diffusion.** Here, we explicitly define the $L_i(q)$ target functions in the loss function $L(q) = \lambda_1 L_1(q) + \lambda_2 L_2(q) + \lambda_3 L_3(q)$. Each target function may also depend on other local information, which we suppress here.

As noted in the main text, $L_1(q)$ is defined by $L_1(q) = |q - \vec{s}|^2$, where $\vec{s}$ is the current vector-valued state of the patch. In the viral dynamics model, each patch takes 5 possible epithelial states and 3 possible endothelial states, leading to 15 possible quantizations, $q$, of an 8 dimensional vector space. Judged by this loss function, the best quantization is the one which is closest to $\vec{s}$ in the $L^2$ norm.

As also noted in the main text, $L_2(q)$ is defined via $L_2(q) = -\log(\vec{c} \cdot q_{epi})$ where $q_{epi}$ is the epithelial portion of the quantization. The vector $\vec{c}$ is the vector of epithelial cell type counts in a Moore neighborhood ($3 \times 3$ square neighborhood) where the count is regularized by adding 1. As the endothelial cells do not clump by type, this loss function ignores them.

The final component of the loss function, $L_3(q)$, assesses the likelihood of the neighborhood under the quantization $q$. To measure this, we collected mean and covariance statistics on all spatial variables in each Moore neighborhood of the $51 \times 51$ grid over 2016 time points (a full run) and 1000 simulations. These spatial variables included both epithelial and endothelial states as well as molecular levels. Then $L_3(q)$ is defined to be, up to an additive constant, the $-\log$ probability under the normal distribution parameterized by these means and covariances. More specifically, $L_3(q) = (\bar{q} - \mu)^T \Sigma^{-1} (\bar{q} - \mu)$ where $\bar{q}$ is $q$ extended by all spatial data in the Moore neighborhood, quantized or not.

Thus, the loss function for our quantization is

$$L(q) = \lambda_1 |q - \vec{s}|^2 - \lambda_2 \log(\vec{c} \cdot q) + \lambda_3 (\bar{q} - \mu)^T \Sigma^{-1} (\bar{q} - \mu).$$

We chose $\lambda_1 = 1.0$, $\lambda_2 = 1.0$, and $\lambda_3 = 0.002$ as this puts the typical values of $L_1$, $L_2$, and $L_3$ at approximately the same order of magnitude and produced reasonable performance.

Quantization naturally introduces an error term $\Delta = \vec{s} - q$ into the global state which must be compensated for elsewhere. The literature contains several algorithms for distributing this quantization error which differ in the order in which we iterate/quantize over the patches and in how the error is distributed at each step. We use the most common algorithm, in which the patches are scanned row-by-row from upper left to lower right. In this iteration scheme, all patches on previous rows and to the left of the current patch have been previously quantized and only patches to the right and on later rows are yet unquantized. See Figure S2, which illustrates the typical case where we distribute the quantization error evenly to all adjacent patches. There are special cases on the right edge of the simulation space and on the bottom row. In these cases, we still distribute quantization error evenly, but only into the subset of the patches indicated in Figure S2 that have not yet been quantized. Several other methods of distributing the quantization error (5, 6) have been explored in the literature, but are largely optimized for visual aesthetics while our method is optimized for locality and conservation of total quantities.

**Sensitivity Analysis for the viral dynamics model.** The viral dynamics model contains a total of 83 distinct parameters for its human model, each with a default value. These parameters are listed and categorized in a separate file. The categories are

- Geometry (2)
- Initial conditions (4)
- Potentially personalizable parameters (59)
- Physical constants (diffusion/evaporation/thresholds) (17)

In the above classification, we viewed a parameter as potentially personalizable if it did not fit into one of the other categories. Throughout, we will fix the parameters considered to be either geometric or a physical constant, leaving 63.

On this parameter space, we performed four varieties of sensitivity analysis. In the first analysis, we performed Latin Hypercube sampling on the parameter space, obtaining 10,000 parameter samples. The model was initialized as specified by the 4 initial condition parameters and run for 2016 time-step under each parameterization, leading to 10,000 model trajectories.

These trajectories were used to fit "global sensitivity matrices". That is, for various values $y$ derived from the trajectories, the corresponding least squares fit $S$ to the equation $y = S[p \quad 1]$ with $p$ the log parameterization. The first considered was log minimum system health vs. log parameterization. In this case, the sensitivity matrix is a column vector and we report it as raw scores in an included file. Here system health refers to the proportion of healthy epithelial cells remaining, as described in (7). In this file, the parameters with the ten largest sensitivities (plus the bias term) are decorated with $\star$. These parameters are listed in Table S2.

Following this, we performed the same analysis, this time fitting the equation $y = S[p \quad 1]$ to $y =$ the system health time-series. In this case, the matrix $S$ is a $2016 \times 64$ matrix and the sensitivity of each parameter corresponds to the $L^2$ column norm. These norms are reported in an included file. In this file, the parameters with the ten largest sensitivities are decorated with †. These parameters are listed in Table S3.

We also considered the sensitivity of the parameters on the full macrostate trajectory. In this case, the matrix $S$ is a $(17 \cdot 2016) \times 64$ matrix where the sensitivity of each parameter corresponds to the $L^2$ column norm. Here, we omit counts of immune cells that are constant in the model as well as the number of healthy epithelial cell, due to a linear dependency. These norms are reported in an included file. In this file, the parameters with the ten largest sensitivities (plus the bias term) are decorated with ‡. These parameters are listed in Table S4.

If we, instead, view the matrix $S$ as a $17 \times 2016 \times 64$ tensor, we can decompose the tensor into time series which indicate which state variables are affected by these parameter variations and at what time. See Figure S4.

We further performed the following procedure for each of the potentially personalizable parameters and parameters determining initial conditions: We fix all other parameters and run the model 2000 times under each 0.8, 0.9, 1.0, 1.1 and 1.2 times the default value of the chosen parameter. For each run, we calculate the minimum value of the system health, that is, the percentage of healthy epithelial cells in the simulation domain.

The resulting empirical cumulative distribution functions (CDF) for the system health is shown in Figure S5. The empirical CDFs for the 0.8, 0.9, 1.1, and 1.2 levels were then compared to the 1.0 level using the 2-sample Kolmogorov-Smirnov test, with results reported in Figure S6. This identified several additional parameters that could be included at the $\alpha = 0.05$ rejection level: `inflammasome_il1_secretion`, `macro_activation_threshold`, `viral_incubation_threshold`, and `infected_epi_t1ifn_secret`.

Combining these lists, the final list of parameters is as shown in Table S5.

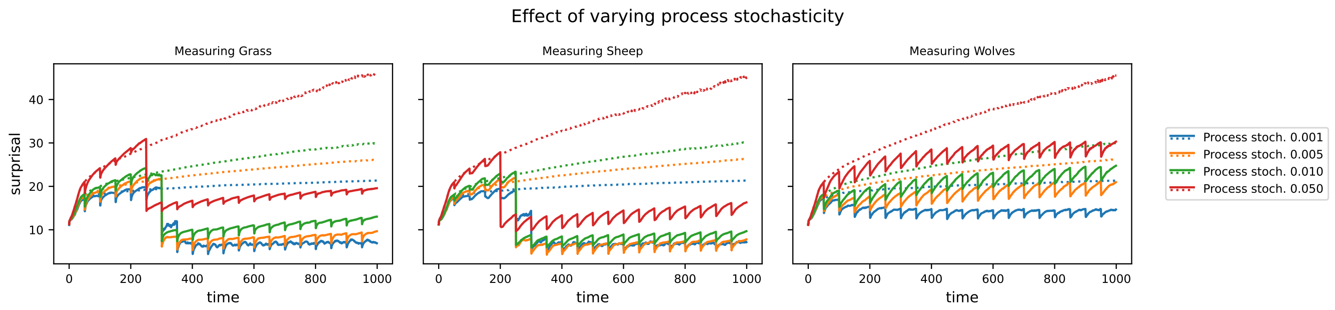


**Fig. S1.** Median surprisal over 1000 runs varying process stochasticity and $R = 0.01$. Dotted lines indicate the trajectory of surprisal when no KF steps are taken and solid lines indicate KF taken every 50 time steps. When the process stochasticity is set too high, the KF cannot compensate for the forgetting aspect of the stochasticity and the surprisal drifts higher after the initial learning period. The $Q = 0.005$ and $Q = 0.001$ levels are small enough for this not to be a problem. The latter performs somewhat better under wolf measurements.

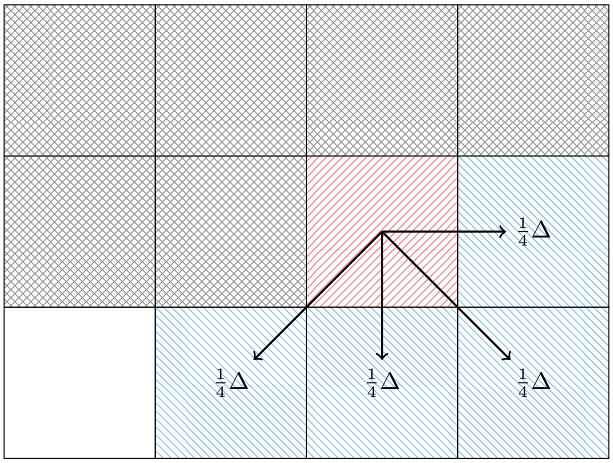


**Fig. S2.** After choosing a quantization for the red patch (north-east hatching), the quantization error $\Delta$ is distributed to the blue patch (north-west hatching). Grey patches (crosshatched) have been previously quantized.

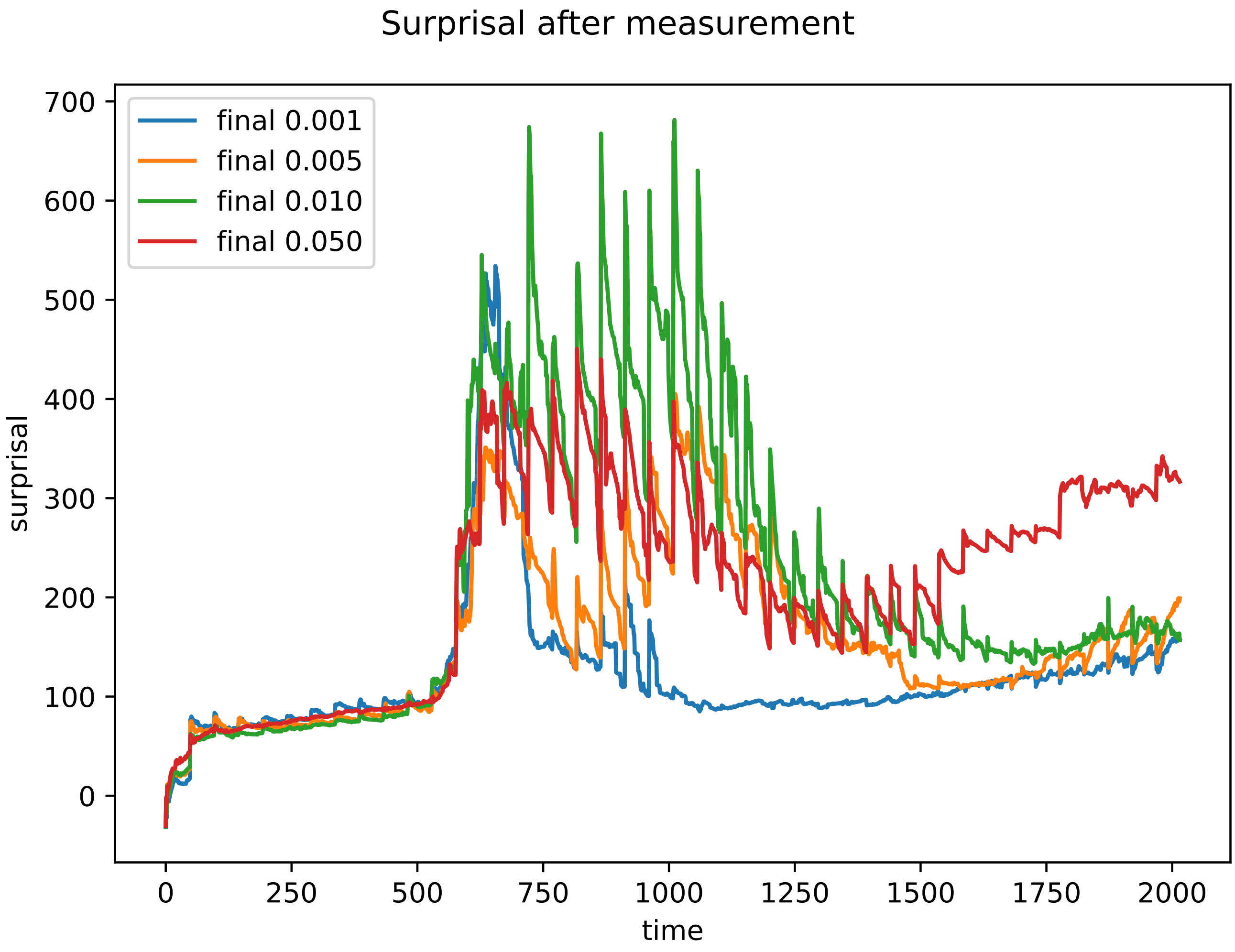


**Fig. S3.** Tuning the process stochasticity for the viral dynamics model. In the spike from $t = 500$ to $t = 750$, the curve with the lowest measurement uncertainty has the highest surprisal due to overfitting by the ABMKF during a period of divergence in trajectories.

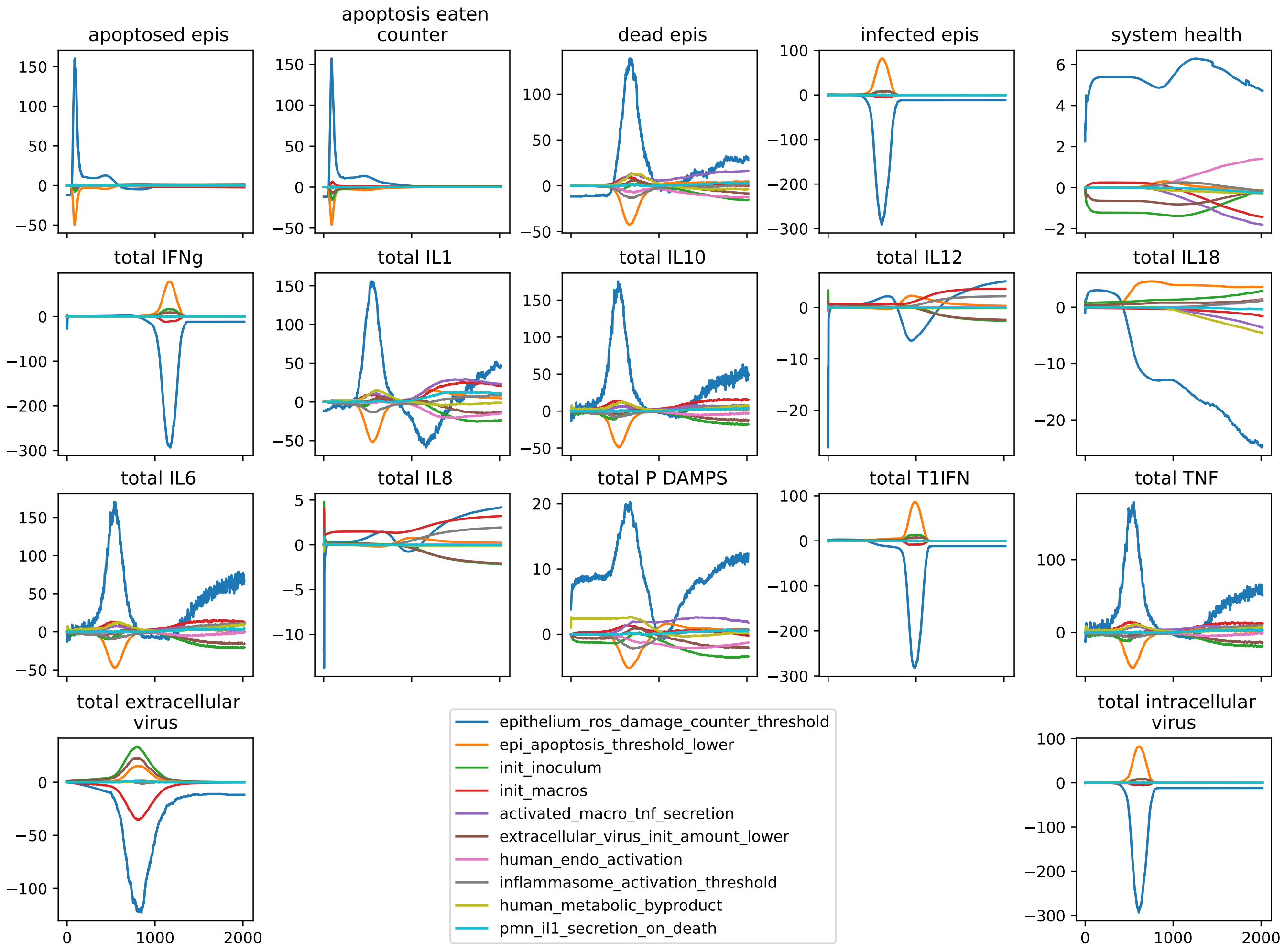


**Fig. S4.** Entries of sensitivity matrix/tensor as time series for ten most sensitive parameters.

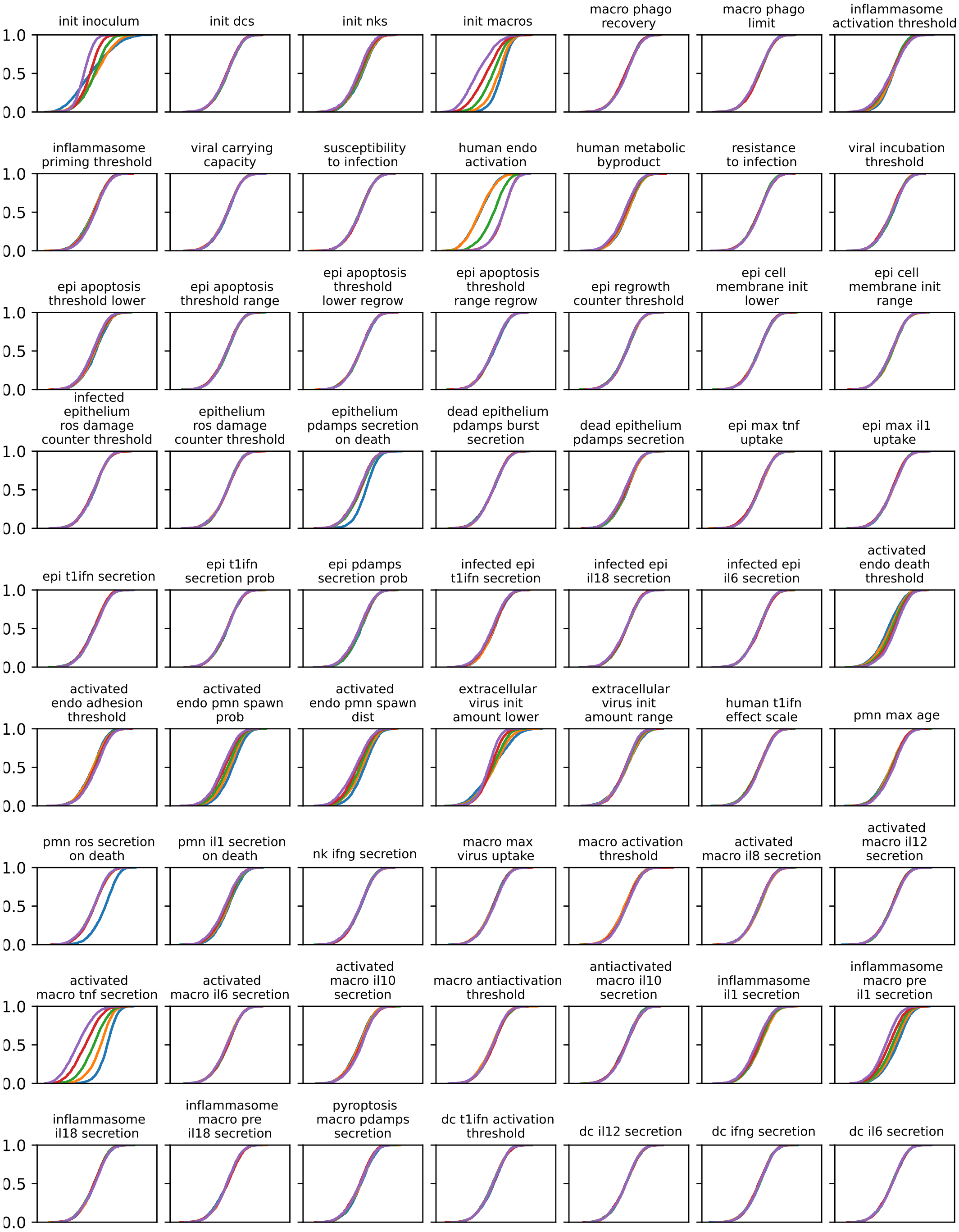


**Fig. S5.** Cumulative distribution (CDF) plots for Kolmogorov-Smirnov sensitivity analysis. Color/levels: Blue/0.8, Orange/0.9, Green/1.0, Red/1.1, Purple/1.2.

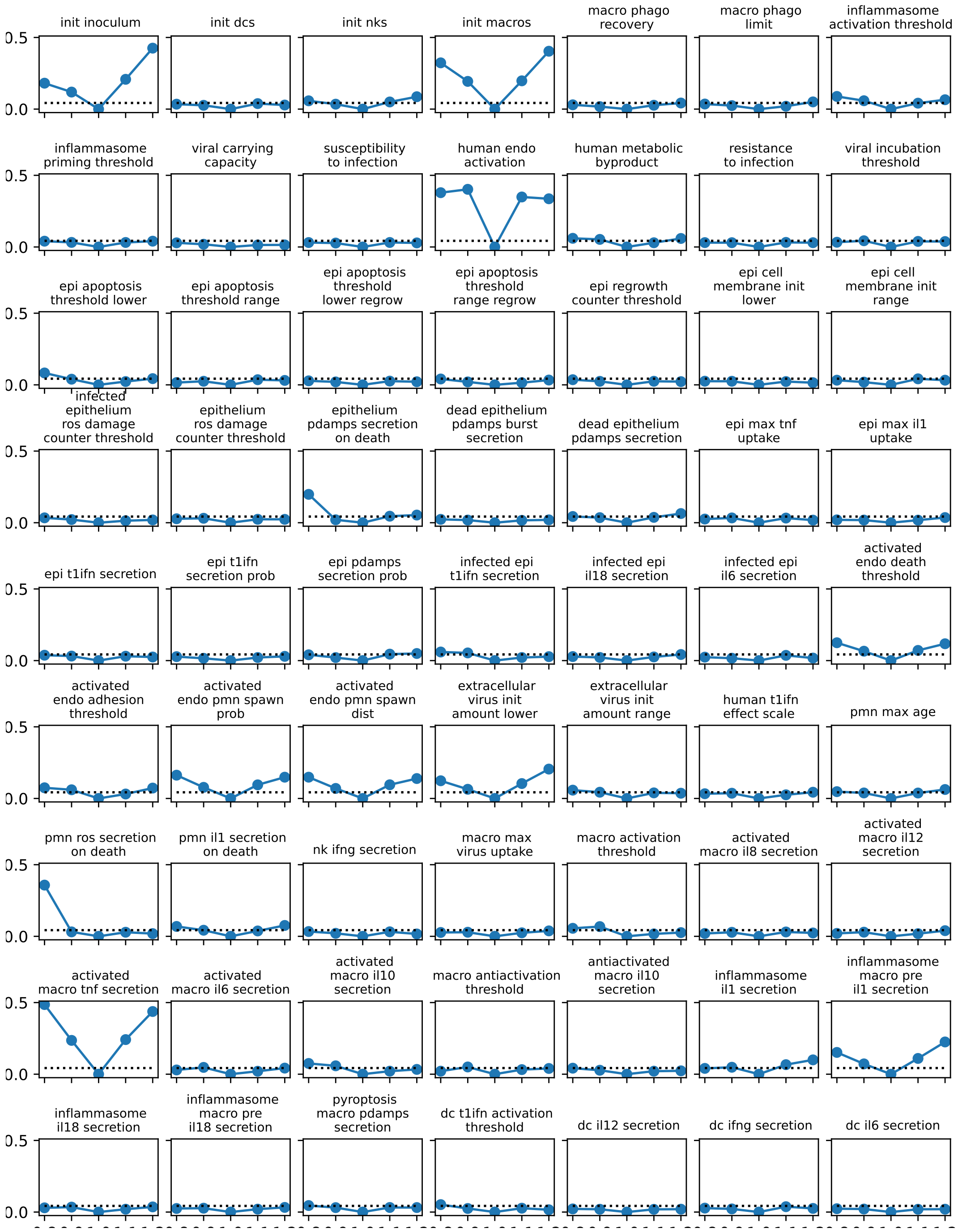


**Fig. S6.** Result of 2-sample Kolmogorov-Smirnov tests comparing the distributions at the 1.0 level with others. Dashed lines indicate the $\alpha = 0.05$ rejection level.

<table>
<tr><th></th><th></th><th>Microstate</th><th>Macrostate</th></tr>
<tr><td rowspan="4">Per-Agent</td><td>Pulmonary Neutrophil</td><td>locations<br>direction<br>age</td><td>count</td></tr>
<tr><td>Macrophage</td><td>locations<br>direction<br>internal virus count<br>activation (bool)<br>cells eaten count<br>virus eaten count<br>pre-IL1<br>pre-IL18<br>pyroptosis counter<br>inflammasome primed (bool)<br>inflammasome active (bool)<br>swollen (bool)</td><td>count<br>apoptosis eaten counter</td></tr>
<tr><td>Natural Killer</td><td>locations<br>direction<br>age</td><td>count</td></tr>
<tr><td>DCs</td><td>locations<br>direction</td><td>count</td></tr>
<tr><td rowspan="3">Spatial</td><td>Epithelium</td><td>state (categorical)<br>ros damage counter<br>regrow counter<br>apoptosis counter<br>intracellular virus<br>cell membrane<br>apoptosis threshold</td><td>state counts<br>total intracellular virus</td></tr>
<tr><td>Endothelium</td><td>activation (bool)<br>adhesion counter</td><td></td></tr>
<tr><td>Molecules</td><td>extracellular virus,<br>P/DAMPS, ROS, PAF, TNF,<br>IL1, IL6, IL8, IL10, IL12,<br>IL18, IFNg, T1IFN</td><td>totals of<br>each</td></tr>
</table>

**Table S1. Micro- and macro-state variables for the viral dynamics model.**

**Table S2. Top ten parameters by $\log$ minimum system health sensitivity.**

| |
|---|
| activated_endo_death_threshold |
| activated_endo_pmn_spawn_dist |
| activated_endo_pmn_spawn_prob |
| activated_macro_tnf_secretion |
| epithelium_pdamps_secretion_on_death |
| epithelium_ros_damage_counter_threshold |
| human_endo_activation |
| inflammasome_macro_pre_il1_secretion |
| init_macros |
| pmn_ros_secretion_on_death |

**Table S3. Top ten parameters by $\log$ system health sensitivity.**

| |
|---|
| activated_endo_pmn_spawn_dist |
| activated_endo_pmn_spawn_prob |
| activated_macro_tnf_secretion |
| epithelium_ros_damage_counter_threshold |
| extracellular_virus_init_amount_lower |
| human_endo_activation |
| inflammasome_macro_pre_il1_secretion |
| init_inoculum |
| init_macros |
| pmn_ros_secretion_on_death |

**Table S4. Top ten parameters by $\log$ full macrostate sensitivity.**

| |
|---|
| activated_macro_tnf_secretion |
| epi_apoptosis_threshold_lower |
| epithelium_ros_damage_counter_threshold |
| extracellular_virus_init_amount_lower |
| human_endo_activation |
| human_metabolic_byproduct |
| inflammasome_activation_threshold |
| init_inoculum |
| init_macros |
| pmn_il1_secretion_on_death |

**Table S5. Combined Parameter list**

| |
|---|
| epithelium_pdamps_secretion_on_death |
| activated_endo_pmn_spawn_prob |
| human_endo_activation |
| activated_macro_tnf_secretion |
| inflammasome_macro_pre_il1_secretion |
| human_metabolic_byproduct |
| pmn_il1_secretion_on_death |
| epi_apoptosis_threshold_lower |
| epithelium_ros_damage_counter_threshold |
| pmn_ros_secretion_on_death |
| activated_endo_death_threshold |
| extracellular_virus_init_amount_lower |
| activated_endo_pmn_spawn_dist |
| inflammasome_activation_threshold |
| inflammasome_il1_secretion |
| macro_activation_threshold |
| viral_incubation_threshold |
| infected_epi_t1ifn_secret |

**Movie S1. Visualization of the Quantization and Error diffusion process in which the number of healthy epithelial cells is increased and number of infected epithelial cells is correspondingly decreased. Initially, the healthy and infected components are scaled to the correct macrostate. Then the Quantization and Error diffusion process scans through the simulation space, adjusting the components to quantized levels while maintaining the macrostate through error diffusion.**

**Movie S2. Second Visualization of the Quantization and Error diffusion process on the epithelial cell patches. Here the model state also includes empty patches as well as apoptosed epithelial cells.**
